# Baseline mutation profiling of 1134 samples of circulating cell-free DNA and blood cells from healthy individuals

**DOI:** 10.1101/089813

**Authors:** Ligang Xia, Zhoufang Li, Bo Zhou, Geng Tian, Lidong Zeng, Hongyu Dai, Fang Deng, Yuancai Xie, Shixin Lu, Xiaohua Li, Chaoyu Liu, Weiren Huang, Jiankui He

**Author notes:** equal contribution (Ligang Xia, Zhoufang Li, Bo Zhou, Geng Tian). Corresponding author: Jiankui He.

## Abstract

The molecular alteration in circulating cell-free DNA (cfDNA) in plasma can reflect the status of the human body in a timely manner. Hence, cfDNA has emerged as important biomarkers in clinical diagnostics, particularly in cancer. However, somatic mutations are also commonly found in healthy individuals, which extensively interfere with the diagnostic results in cancer. This study was designed to examine the background somatic mutations in white blood cells (WBC) and cfDNA for healthy controls based on the sequencing data from 1134 samples, to understand the patterns and origin of mutations detected in cfDNA. We determined the mutation frequencies in both the WBC and cfDNA groups of the samples by a panel of 50 cancer-associated genes which covered 20K nucleotide regions using ultra-deep sequencing with average depth >40000 folds. Our results showed that most of mutations in cfDNA originated from WBC. We also observed that NPM1 gene was the most frequently mutant gene in both WBC and cfDNA. Our study highlighted the importance of sequencing both cfDNA and WBC, to improve the sensitivity and accuracy for calling cancer-related mutations from circulating tumor DNA, and shielded light on developing the early cancer diagnosis by cfDNA sequencing.

## Introduction

A high percentage of cancers are diagnosed at advanced stages, resulting in the failure of curative treatment and poor life quality for patients (*1, 2*). Cell-free DNA (cfDNA) in the blood circulation system of tumor patients and circulating tumor DNA (ctDNA) in the plasma have emerged as potential important biomarkers for cancer monitoring and treatment (*3*). Recently, multiple groups had reported that levels of plasma ctDNA increased in cancers (*4, 5*). Some groups also proposed that cfDNA could be used as a noninvasive, sensitive tool for cancer early diagnosis.

According to what has been elucidated, the circulating cfDNA is a mixture of DNA from blood cells, virus, solid organs and many other sources. More than 90% of cfDNA is from the debris of blood cells in healthy individuals. However, tumor-related ctDNA is present in small amount of 0.1%-0.01% in the plasma cfDNA, which means accurate detection of ctDNA for cancer early diagnosis must remove the background of somatic mutations originated from the blood cells (*6–8*). However, there is limited knowledge available for the somatic mutations in normal cells, particularly in white blood cells that are the main contributor of the background. It is reported that human genomic mutation happened at rate of ∼2.5*10^-8^ per base per cell generation (*9*). Campbell group studied the background mutation spectrum of skin samples(*10*). They focused on 74 cancer genes in normal skin samples from four individuals, proving somatic mutations accumulate in normal cells, although the mechanisms were poorly understood. In a word, the baseline spectrum of somatic mutations in healthy individuals is urgently required before the detection of ctDNA can be useful in early cancer diagnosis.

With the dramatically decreased cost of next generation sequencing in recent years, it is now possible to screen a large number of individuals at ultra-deep sequencing to identify cancer related mutations (*11–15*). Development of molecule technology to suppress the technique noise also makes it possible to pinpoint the mutations, fusions and copy number variations related to cancer in the ultra-low concentration of ctDNA. Both academic researcher groups and industry players are chasing the pan-cancer screening by a simple blood draw. However, the reliable and accurate application of ctDNA detection needs a better understanding of background somatic mutation information in healthy individuals,

Our studies were designed to reveal the background somatic mutations in cfDNA by ultra-deep sequencing for 50 cancer-associated genes in 1134 samples, providing the critical information of baseline profile of somatic mutations in healthy individuals, which will fill the gap in early cancer diagnosis and personalized cancer therapy.

## Material and methods

### Patients

This study was approved by the declaration of the Second People’s Hospital of Shenzhen (The First Affiliated Hospital of Shenzhen and the other sources). A written consent was obtained from each participant and all analyses were performed anonymously. Totally 1134 samples were included in this study (**Table S1**), including 820 cfDNA samples and 314 white blood cell samples. We have 56.6% females and 43.4% males. The age of samples ranges from 16 to 86, with the median 42 years old. There is no significantly genetic diverse in the samples based on our analysis of clinical information. The statistical analysis of sample information is summarized in **Table 1**

**Table 1.** Summary of patient information

| Age years | Parameter value |
| --- | --- |
| Mean (SD) | 43.1 ( 12.8 ) |
| Median (range) | 42.5 ( 16 - 86 ) |
| 10 - 19 n ( % ) | 2 ( 0.2 ) |
| 20 - 29 n ( % ) | 128 ( 15.6 ) |
| 30 - 39 n ( % ) | 233 ( 28.4 ) |
| 40 - 49 n ( % ) | 194 ( 23.7 ) |
| 50 - 59 n ( % ) | 158 ( 19.3 ) |
| 60 - 69 n ( % ) | 87 ( 10.6 ) |
| 70 - 79 n ( % ) | 17 ( 2.1 ) |
| 80 - 89 n ( % ) | 1 ( 0.1 ) |
| Sexr n ( % ) |  |
| Female n ( % ) | 464 ( 56.6 ) |
| Male n ( % ) | 356 ( 43.4 ) |
| smoking n ( % ) |  |
| Female n ( % ) | 62 ( 7.6 ) |
| Male n ( % ) | 149 ( 18.1 ) |
| Family tumor history n ( % ) |  |
| single parent n ( % ) | 81 ( 9.9 ) |
| both parents n ( % ) | 2 ( 0.2 ) |

### Blood plasma isolation

For each sample, 10 ml of blood in a cell free DNA collection tube (VanGenes, Xiamen, China) was collected, then centrifuged at 1600 g for 10 minutes at 4^o^C to roughly separate the whole into plasma and blood cells. The upper phase was then transferred into a new tube, leaving around 8mm of “buffering” layer from the buffy coat after the centrifuge and avoiding contaminating the plasma layer by blood or cell debris. The plasma was further centrifuged at 16000g for another 10 minutes to remove the left cell debris. The upper clear layer was then aliquoted into 2ml tubes and clearly labeled with the patients’ identity and immediately stored at −80°C for cfDNA extraction. Each extraction of cfDNA was from 1ml of plasma stored here.

### DNA extraction

cfDNA was extracted using E.Z.N.A Circulating DNA Kit (Omega Bio-tek, GA, USA). Genomic DNA of white blood cells was extracted by Qiagen DNA mini kit (Qiagen, Hilden, Germany). The concentration of extracted DNA was measured using the Qubit 2.0, dsDNA high sensitivity assay (Life Technologies, Carlsbad, CA).

### Multiplex PCR and sequencing library construction

We developed a multiplex PCR method to enrich cancer-associated genes. Totally 50 cancer-associated genes were included which were amplified by 207 amplicons and covered 22027bp-region. 2800 mutations are defined as the hotspot mutations according to Mayo Clinic. The 50 genes listed in Table S2. 10 ng of DNA per sample was used for amplicon production by multiplex PCR. The resulting multiplex PCR reaction pool was further used in sequencing library preparation using the reagents provided in the same kit following the manual’s instructions. The sequencing library was then sent to WuxiNextCODE on the illumina Hiseq X10 platform (illumina, Beghelli, CA, USA)

### Data filtering and analysis

We performed ultra-deep target sequencing on 50 cancer-associated genes for both cfDNA and blood-cell DNA. For each cfDNA and WBC sample, the average sequencing depth is 47218X and 54284X. The sequencing data was first mapped to human reference genome (human genome build19; hg19) by Burrows-Wheeler transformation (BWA, Version: 0.7.5a-r405) software package, converted to mpileup format for downstream analysis. We set two filtering criteria to filter the reads: 1). Reads sequences with mutation rate higher than 5% in a single reads were deleted. 2). Bases with base quality lower than 30 were deleted. Loci with sequencing depth less than 10000x were removed for further analysis.

### Mutation rate in cfDNA and white blood cells in the population and in individuals

We analyzed the correlation of mutation rate between cfDNA and genomic DNA of white blood cells. The mutation rate is defined as mutant allele frequency divided by total reads covering the locus. For example, for a particular position, the total sequencing depth is 10000, we obtain 9990 with A, 3 with C, 5 with G, 2 with T, then the mutation rate for this position is (3+5+2)/10000=1/1000. Among 1134 samples, we only use these 309 pair-of samples of which both cfDNA and white blood cells are available from the same person (collected at the same time). Totally 22027 nucleotide loci in 50 gene were included in this study. We calculated the average mutation rate of each position in 309 individuals in white blood cells and cfDNA respectively. We removed the positions with mutation rate high than 0.1.

### Reproducibility evaluation

We draw blood from an individual, spilt it into two halves, and performed multiplex PCR and sequencing independently, to validate the reproducibility of our methods. By comparing the sequencing data of two replicates in WBC samples, we were able to evaluate the stability of the methods and remove background noises. Positions with sequencing depth lower than 10000x and mutation rate less than 0.003 is excluded in the correlation study.

## Results

### The mutations detected in cfDNA are highly correlated with mutations in white blood cells

We performed ultra-deep target sequencing on 50 cancer-associated genes for both plasma DNA (cfDNA) and blood-cell DNA (**Supplementary Figure S1**). We analyzed the correlation of mutation rate in 309 WBC samples and the corresponding cfDNA respectively. As shown in Fig. 1a, each dot represents the average mutation rate of one position in both WBC and cfDNA. Pearson’s correlation study suggested that cfDNA and white blood cell is highly correlated (adjusted R^2^=0.92) **(Figure 1a, and Supplementary Figure S2a-c**). We plot the data from only one individual, the correlation of the WBC VS cfDNA in one individual sample is also highly correlated, with R^2^ =0.80 **(Figure 1b)**. The majority of mutations in cfDNA were contributed by somatic mutations in the blood cells. These results also highlighted the importance of sequencing both cfDNA and blood cells, to remove the background mutations contributed by blood cells.

**Figure 1.**
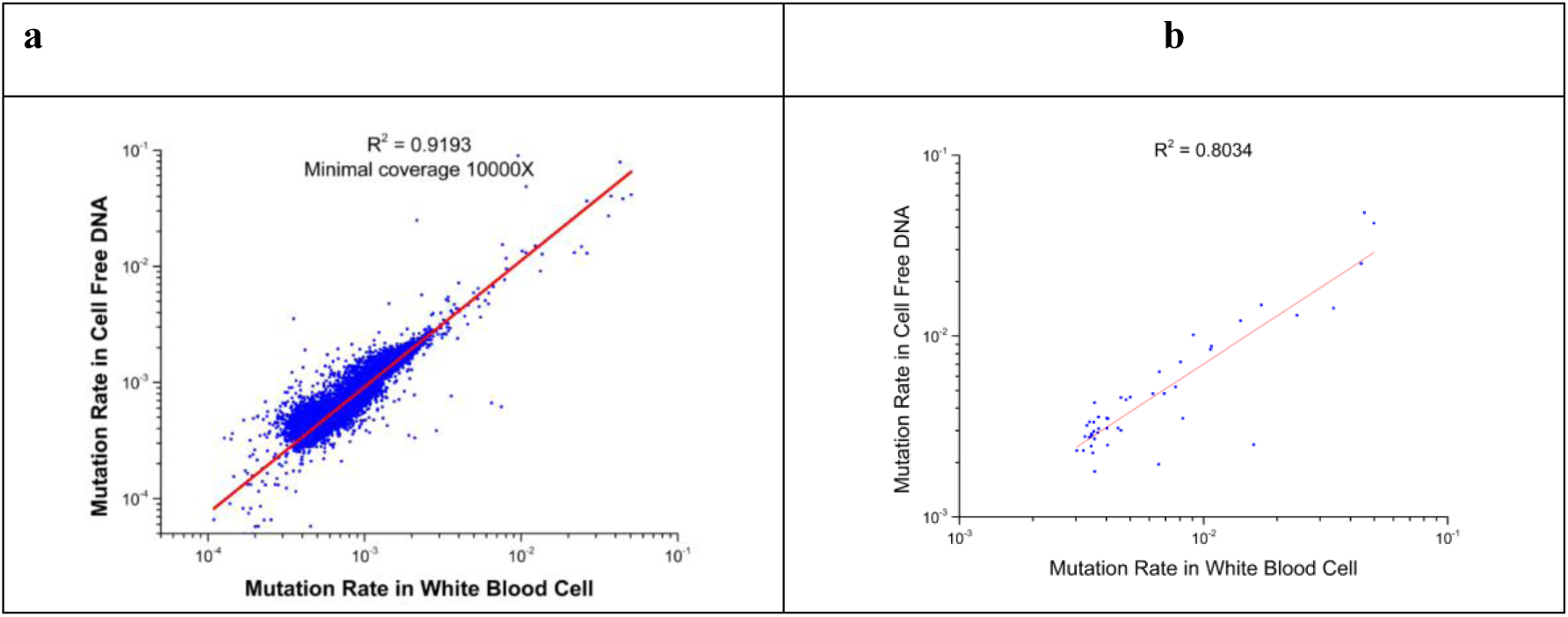
Correlation study of mutation rate in ctDNA and white blood cells. (a) We analyzed the correlation of mutation rate between cfDNA and genomic DNA of white blood cells. 309 paired-samples are included in this study. Each dot represents the mutation rate of one position in WBC and its corresponding cfDNA, averaged for 309 samples. (b) Correlation study of one individual. Here, only the total sequencing depth larger than 10000x and mutation rate larger than 0.3% in both WBC and cfDNA are selected in this analysis.

### Reproducibility

One of the key issues to be addressed is to distinguish the true biological variations from experimental noise or errors. Two technical replicates were performed for both WBC and cfDNA samples from the same individual. We compared the mutation rate in two WBC samples or two cfDNA samples. The reproducibility validation experiment in two WBC samples was used to estimate the PCR error rate and sequencing errors in our study cohort. Our data showed that the mutation rate in WBC is highly correlated (R^2^=0.9672), which suggested the mutations with frequency >0.3% were highly reproducible **(Figure 2a)**. The two cfDNA samples have good correlations (R^2^=0.85), but were not as good as WBC samples **(Figure 2b)**. The lower correlation of cfDNA replicates may arise from both biological and technique variations. One of the possible reasons was that the cfDNA is more complex in compositions. cfDNA may from debris of different tissues and each subtype may contain very small amount DNA. The Limit amount of starting material of cfDNA for experiments can also introduce technique variations during the amplification process.

**Figure 2.**
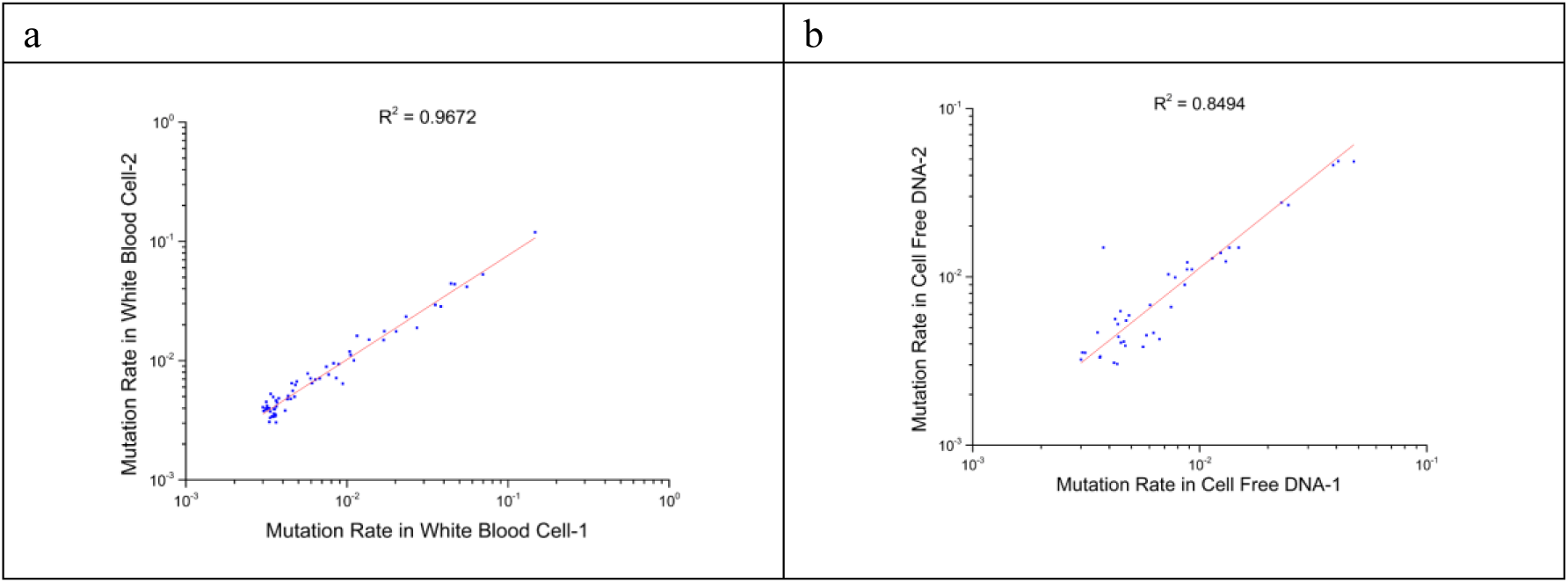
Reproducibility validation. We compared the sequencing information of two repeated WBC samples (a) and cfDNA samples (b) to evaluate the reproducibility of the methods. Only the total sequencing depth larger than 10000x and mutation rate larger than 3/1000 are included in the correlation study.

### NPM1 is the most frequently mutated gene

The mutation frequency of 50 genes were analyzed and ranked by comparing 309 cfDNA samples with average reads depth more than 10000X (**Figure 3a**). The top 4 mutated genes were NPM1, PI3K, KIT and PTEN (**Figure 3b** and **Supplementary figure S3 b-d**). The average mutation rate of NPM1 gene in each nucleotide was up to 0.12%. This feature was observed in both the cfDNA and WBC (**Supplementary figure S3a**), suggesting that the major origin of NPM1 mutations was from blood cells. These results suggest that cautions need to be taken for these blood prone mutations when using the cfDNA as the diagnostic marker.

**Figure 3.**
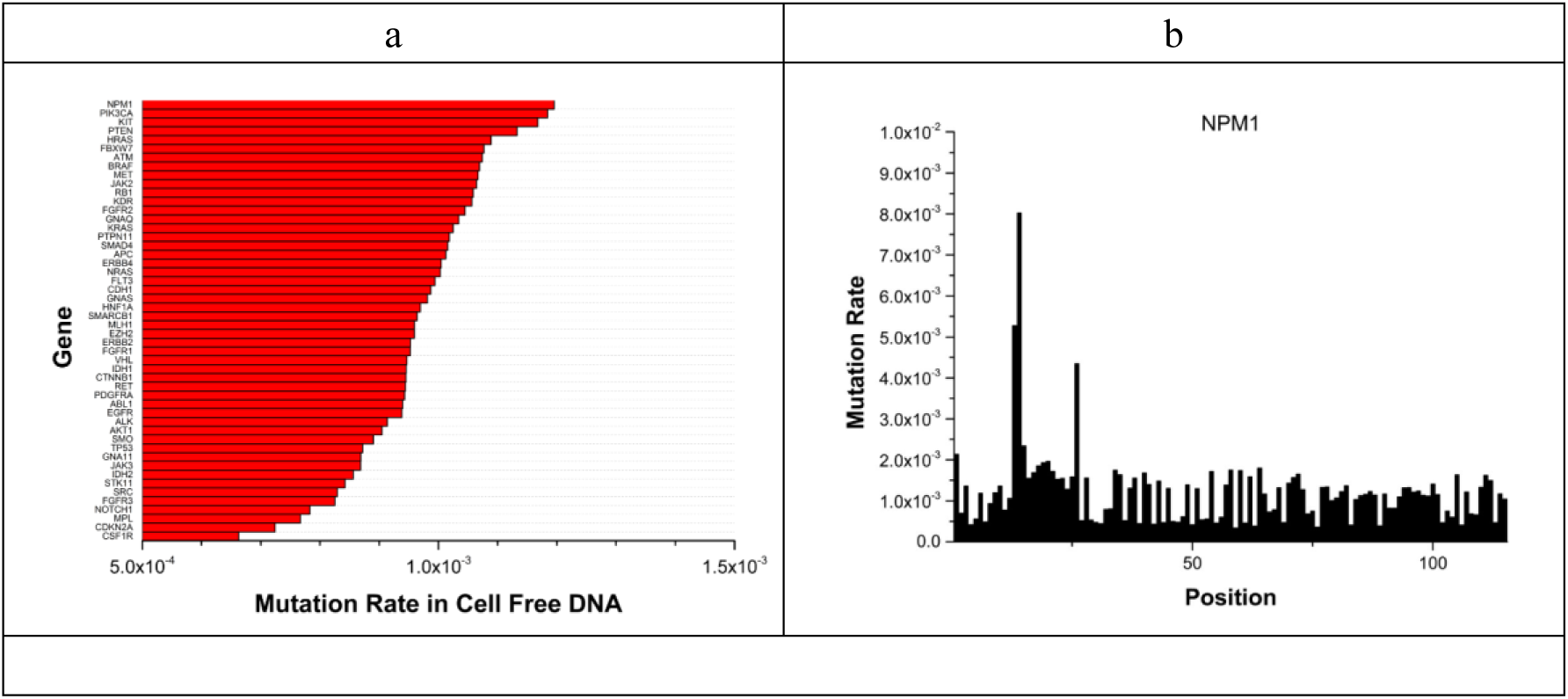
Ranking of average mutation rate of 50 cancer associated-genes. (a) Mutation rate per position for 50 genes. (b) The mutation rate of each position in NPM1 gene. Here, only one amplicon was designed for NPM1 in our study.

## Discussion and conclusions

In recently years, cfDNA and ctDNA has emerged as the research frontier of non-invasive cancer biomarker for the detection, monitoring and treatment of cancer (*16–21*). However, there were several challenges before cfDNA and ctDNA can be widely in cancer early diagnosis The first challenge is the system errors derived both from the PCR and the sequencing. ctDNAs are released to the peripheral blood in the very beginning of tumor initiation, far before the tumor can be detected by either CT or other clinical tools. However, in very early stage of tumor, the amount of ctDNA is as low as 0.1-0.01%. The sequencing alone may introduce ∼0.1% error by Illumina sequencing chemistry. Several groups have proposed to use digital barcoding of every molecule to reduce the errors of PCR and sequencing (*22*). We also tried digital barcoding and the results will be published in the following articles.

The second challenge is distinguish tumor mutations from background mutations from WBC. Since more than 90% of cfDNA are from blood cells, we also observed that mutations in cfDNA were highly correlated to that in the WBC. Therefore, the majority mutations detected in cfDNA were the background mutations from WBC. The cfDNA also contained many other sources of DNA and its composition and origin varies by individual and by different disease states. It is critical to perform systematic analysis of WBC somatic mutations, and investigate the origin of cfDNA to understand all factors that may contribute the mutation spectrum of cfDNA. In our study, we demonstrated the importance of sequencing both cfDNA and WBC for every individual.

Somatic mutations in healthy group are very prevalent, with average mutation number around 2 to 6 per 1 million bases (*10, 22, 23*). There is little information available to determine these biologically significant variations in cfDNA. This study revealed the baseline mutation spectrum in WBC and cfDNA of healthy groups and can somehow fill the gaps for the early cancer diagnosis. We observed an interesting pattern in WBC that NPM1 gene is mutated in a very high rate. The average mutation rate of all 1134 samples is as high as 0.12%. The NPM1 mutations behave as gatekeepers for leukemia and have been reported to be important cancer gene in multiple type of leukemia(*24*). It was quite a surprise that TP53 gene was not the most frequent mutant gene in cfDNA samples. This observation can be explained by fact that TP53 mutations were more frequent in solid organ tumors, while the cfDNA are mainly from blood cells. Therefore, we need to calibrate the background mutations in WBC and cfDNA of healthy individuals carefully to reduce false positive rate during early cancer diagnosis.

Our study can help defining the threshold of mutation detection of ctDNA by removing the background mutations in WBC and cfDNA of healthy population. In the future, three aspects should be further improved: 1) Larger sample size is required to refine the baseline spectrum. 2) Experimental methods that can increase the detection limit and remove technique noise are highly desired. 3) Efficient bioinformatics pipeline need to be developed to remove the noise and increase the accuracy and sensitivity in mutation detection of ctDNA.

